# Auditory Figure-Ground Segregation is Impaired by High Visual Load

**DOI:** 10.1101/216846

**Authors:** Katharine Molloy, Nilli Lavie, Maria Chait

**Affiliations:** Institute of Cognitive Neuroscience, University College London, London WC1N 3AR, United Kingdom; Ear Institute, University College London, London WC1X 8EE, United Kingdom

**Keywords:** auditory scene analysis, MEG, Attention, Load Theory;

## Abstract

Figure-ground segregation is fundamental to listening in complex acoustic environments. An ongoing debate pertains to whether segregation requires attention or is ‘automatic’ and pre-attentive. In this magnetoencephalography (MEG) study we tested a prediction derived from Load Theory of attention^1^ that segregation requires attention, but can benefit from the automatic allocation of any ‘leftover’ capacity under low load. Complex auditory scenes were modelled with Stochastic Figure Ground stimuli (SFG^2^) which occasionally contained repeated frequency component ‘figures’. Naive human subjects passively listened to these signals while performing a visual attention task of either low or high load. Whilst clear figure-related neural responses were observed under conditions of low load, high visual load essentially abolished the neural response to the figure in auditory cortex (Planum Temporale, Heschl’s gyrus). We conclude that fundamental figure-ground segregation in hearing is not automatic but draws on shared resources across vision and audition.

## Introduction

Figure-ground segregation – the process by which an auditory object is perceptually extracted from the aggregate sound input – underlies key aspects of listeners’ ability to make sense of the acoustic environment, including recognizing individual sounds within crowded scenes or understanding speech in noise. The extent to which segregation depends on attention has been a long standing question in hearing research,^3–6^ crucial for establishing how the acoustic environment is represented when attention is directed away from sound. However, despite decades of research, the answer has remained elusive.

Previous studies^7–11^ have predominantly focused on mixtures of signals that are separated in frequency, based on the understanding that segregation occurs when spatially distinct populations of neurons are activated in the primary auditory cortex (for reviews see ^5, 12, 13–19^). However, natural sounds rarely exhibit the frequency separation used in these standard stimuli, suggesting that they may not adequately capture the challenges faced by the auditory system in complex acoustic environments.

Recent work suggests that in addition to frequency separation, another cue for the perceptual organization of acoustic scenes is ‘temporal coherence’ ^2, 4, 20–23^. Sounds with similar temporal properties are bound into one perceptual object, via a process that detects correlations between the statistics of individual frequency channels in auditory cortex^2, 4, 24^. This process provides segregation cues despite spectral overlap between auditory objects, and can thus play a vital role in segregating the spectrally broad, dynamic sounds we experience in natural environments.

The Stochastic-Figure ground stimulus (SFG^2, 25, 26^; Figure 1) has been proposed as a means by which to model complex scene analysis. Signals are made of multiple, rapidly changing, random frequency components, which form a stochastic background. ‘Figures’ are introduced by repeating a small subset of components throughout the signal, resulting in the percept of a bound auditory object that is segregated from the random ground^2^. Importantly, similarly to natural sound mixtures, and in contrast to most previous work, the figure is not dissociable from the background based on instantaneous cues and can only be identified by integrating information over both frequency and time.

**FIGURE 1:**
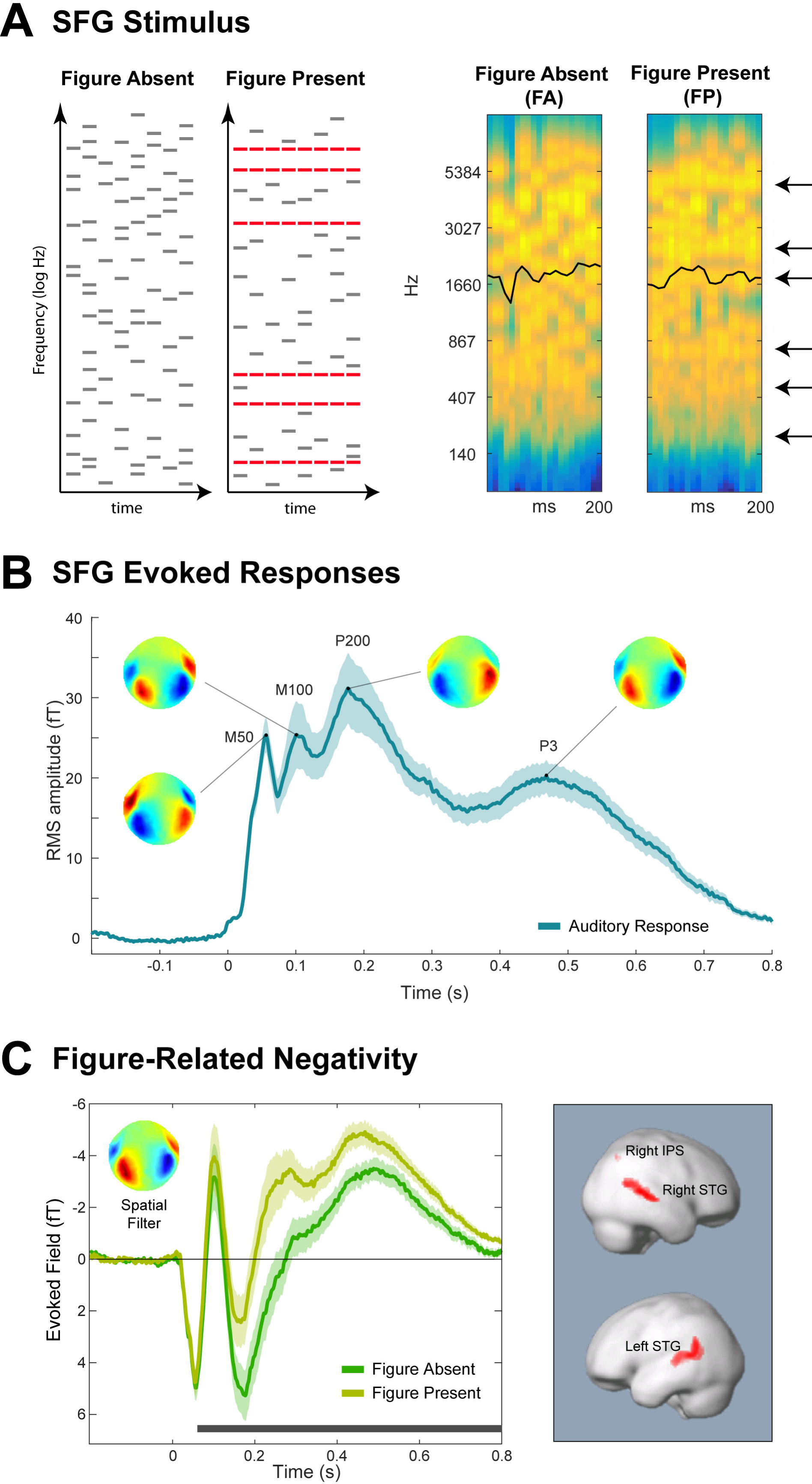
Stimuli and Basic Meg Responses Under Low Load (Experiment 1) **A. Example schematics (left) and spectrograms (right) of the Stochastic Figure-Ground (SFG) stimuli**. Stimuli, adapted from Teki et al.^2, 26^, consisted of a succession of chords, each comprised of multiple frequency components. The *‘figure-absent’* (FA) stimuli were chords formed of random frequencies, while the *‘figure-present’* (FP) stimuli were constrained so that some of the frequencies (indicated in red in the schematic representation and by black arrows in the spectrogram) were repeated in each chord of the stimulus, producing an auditory ‘figure’. The spectrograms were generated with a filterbank of 1/ERB wide channels (Equivalent Rectangular Bandwidth^63^) equally spaced on a scale of ERB-rate. Channels are smoothed to obtain a temporal resolution similar to the Equivalent Rectangular Duration^64^. This models processing in the auditory periphery to produce a representation of the stimulus similar to that available to the central nervous systems. Black lines overlaid on the spectrograms indicate the ERB-weighted spectral centroid. **B. Overall response to the SFG stimuli (collapsed over FA/FP conditions) under low load (Experiment 1)**. Plotted is the mean RMS (instantaneous power) of stimulus-evoked activity collapsed over stimulus conditions. Associated scalp topographies at major peaks are provided. Error bars reflect the standard deviation of bootstrap resamplings. **C. Figure-related negativity (Experiment 1)** Evoked fields calculated using a spatial filter which maximised the difference between FP and FA responses (inset – see methods). Evoked fields in this and subsequent figures are plotted with the M100 as an ‘upwards’ peak, to match the convention used in EEG for its counterpart, the N1. Error bars reflect the SD over bootstrap resamplings for each condition, and significant differences between the conditions are marked by horizontal black bars at the bottom of the plot. Right panel shows source-level contrast: regions where FP trials showed greater activity than FA trials are indicated in red. No regions were found to be significant in the opposite direction to that displayed.

Accumulating work has shown that listeners are highly sensitive to SFG figures; brain responses to the emergence of the figure have been consistently observed in naïve, distracted, listeners performing an incidental task^24–26^. This has been interpreted as indicating an automatic (pre-attentive) segregation process. However, it is well established that attention is only fully withdrawn from a stimulus when a separate, prioritized task involves a high level of perceptual load that exhausts capacity (e.g.^1, 27, 28^). The incidental tasks used previously, where participants either listened passively or watched a subtitled film, are unlikely to have placed a high enough instantaneous demand on processing to ensure that attention was diverted from the auditory stimuli. Thus, it remains possible that figure ground segregation does depend on attention and cannot occur when attention is fully withdrawn from the sound. Investigating the impact of perceptual load on the neural response to SFG signals can thus provide a crucial window into understanding the extent to which analysis of spectro-temporally complex auditory signals draws on attentional resources.

Here we actively divert attention away from the auditory stimuli using a visual task of high or low perceptual load. We show that load has a significant effect on segregation such that under conditions of high visual load, the neural response to the auditory figure is essentially abolished. This indicates that in fact, identification of the figure does depend on attention, and that even basic auditory scene analysis draws on processing resources which are shared across the visual and auditory senses.

## Results

### Experiment 1: MEG to short SFG sequences

In this series of experiments, we focus on brief (~200ms long) SFG bursts that occasionally (in 50% of the trials) contain a figure (Figure 1A). As a first step, Experiment 1 was designed to characterize the MEG response under passive listening conditions. To maintain vigilance, participants were engaged by a simple, very low load, incidental, visual task.

Figure 1B shows the evoked response to the SFG stimulus, collapsed over figure-present and figure-absent trials. Because these data are comprised of 20 components (i.e. reflect the independent activity of many processes), their dynamics are summarized by calculating instantaneous power (RMS; see methods) across channels. Visible is the characteristic succession of onset peaks (M50, M100, P200 at 50, 100 and 200 ms post onset), followed by a P3 response from ~300-700 ms post stimulus onset.

A source separation analysis (see methods) was used to identify neural activity which is most different between figure-present (FP) and figure-absent (FA) trials. The associated spatial filter (in the inset) was applied to the data to produce the time series in Figure 1C. The response to the FP trials relative to FA trials is characterized by a sustained ‘negativity’ (i.e. in the same direction as the M100 peak) which was significant from 60ms post stimulus onset and persisted throughout the rest of the trial. We refer to this effect as the ‘figure-related negativity’ (FRN). The pattern is generally reminiscent of the object-related negativity (ORN) response which has been observed when simultaneous auditory stimuli are perceived as two objects rather than one (usually a mistuned harmonic within an otherwise harmonic chord^29–32^, and more recently also in figure-ground stimuli similar to those used here^33^). The ORN is typically superimposed on the N1-P2 complex, peaks between 150-300ms post stimulus onset and can occur even when auditory stimuli are not actively attended.

These results confirm that there is a measurable neural response to the presence of the figure even during very brief SFG signals, consistent with previous behavioural reports^2^, and despite the fact that the sounds were not explicitly attended. The fact that a response to the figure can be seen within 60ms of scene onset (just over 2 chords), suggests a very rapid, sensitive figure-ground segregation process.

Source localisation revealed several brain regions where activity differed significantly between FP and FA trials (Fig 1C, Table 1). FP stimuli showed greater activity in bilateral superior temporal gyri and right superior and inferior parietal lobules. This activity is consistent with the findings of Teki et al.^25, 26^ that the SFG stimuli evoke figure-specific activity along the superior temporal planes, superior temporal sulci, and also within the intraparietal sulci.

**Table 1:** Experiment 1, Effect of Figure. Source estimates for the difference between FP and FA trials. * indicates a small volume correction.

| Cluster | *p*(FWE-corr) | Peaks |  | Co-ordinates (x,y,z) |  |  |
| --- | --- | --- | --- | --- | --- | --- |
|  |  | *t* | *p*(uncorr) |  |  |  |
| Cortical Structures |  |  |  |  |  |  |
| **Left Temporal Lobe:** | <.001 | 5.12 | <.001 | -58 | -42 | 10 |
| Superior Temporal Gyrus, BA 22 |  | 4.63 | <.001 | -58 | -52 | 24 |
| **Right Temporal Lobe:** | <.001 | 5.72 | <.001 | 56 | -42 | 10 |
| Superior Temporal Gyrus, BA 22 |  |  |  |  |  |  |
| **Right Parietal Lobe:** | 0.026* | 3.54 | <.001 | 34 | -66 | 46 |
| Superior Parietal Lobule, Inferior Parietal Lobule, BA 7 |  | 3.32 | 0.001 | 34 | -62 | 48 |

### Experiment 2: Effect of Perceptual Load on Figure-Ground Segregation

To understand how figure-ground segregation is affected by the availability of processing resources, we recorded MEG responses to non-attended SFG signals while attention was engaged by a concurrently presented visual task which placed different levels of load on perceptual processing. The auditory stimuli were identical to those in Experiment 1, but with fixed parameters: coherence 6, chord duration 25ms and chord number 8, producing a 200ms long stimulus. A feature vs. combination visual search task was used to implement different levels of visual perceptual load^1, 34^. Figure 2B shows a schematic diagram of the trial structure. The search array, presented for 200 ms, consisted of five coloured shapes spaced around a (non-visible) circle centred at fixation. The low load (LL) task, colour search, required participants to respond to any blue shape. The high load (HL) task, colour-shape combination search, required participants to respond to a red circle or a green square. On half of the trials the visual search array was accompanied by an auditory stimulus. Participants were told that the sounds should be ignored and were encouraged to focus on the visual task.

**FIGURE 2:**
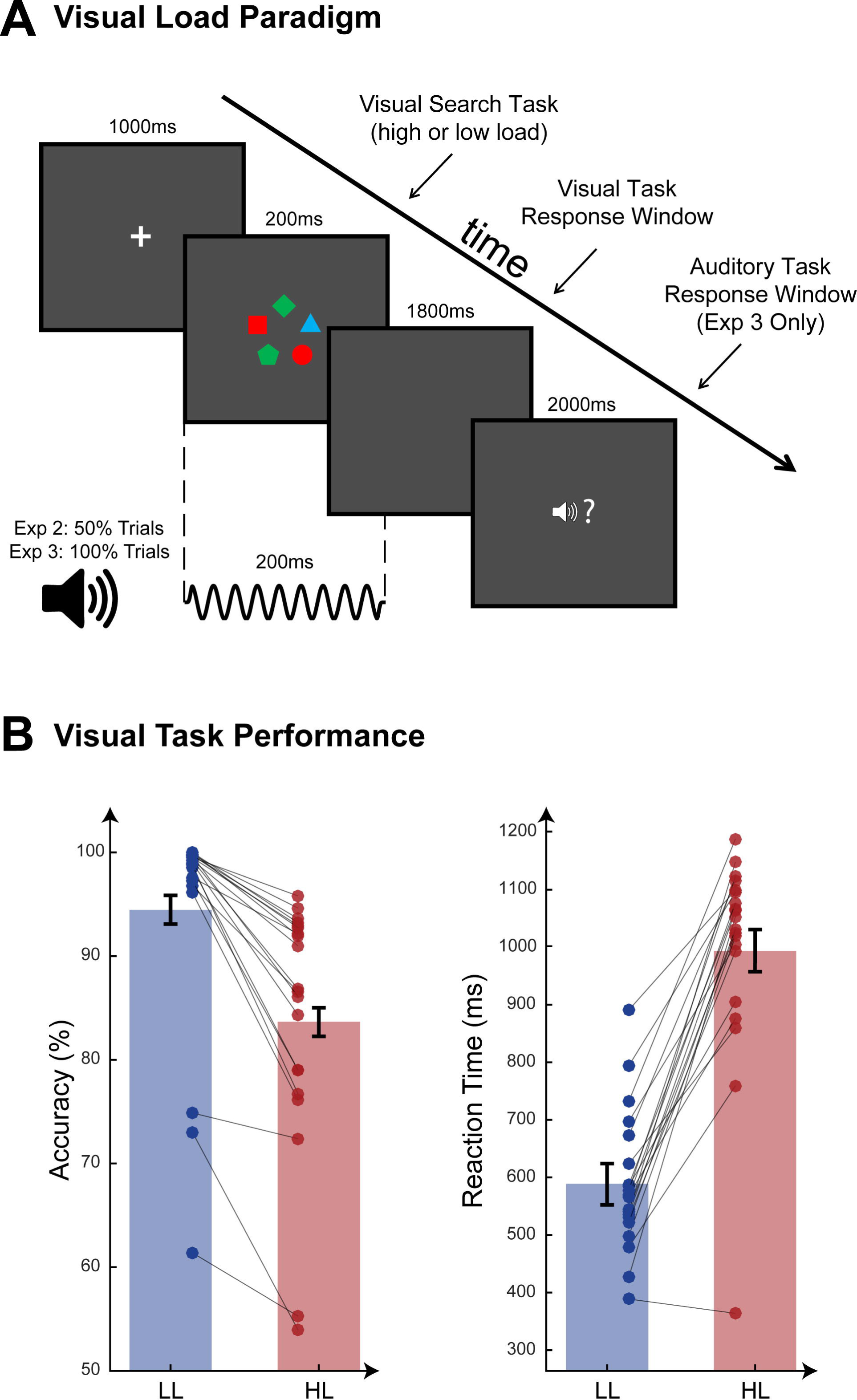
Experiment 2 - Visual Load Task. **A. Load Task Paradigm used in Experiments 2 and 3**. Low load (LL) task was a colour feature search; high load (HL) task was a colour-shape combination search. Auditory stimuli occurred on 50% of trials in Experiment 2 (MEG) and 100% of trials in Experiment 3 (Behavioural Dual Task). When present, auditory stimuli occurred at precisely the same time as the visual search array. The response window for the auditory target was displayed only during Experiment 3 when there was an active auditory task. **B. Visual task behavioural data from Experiment 2 (MEG).** Mean values for accuracy (percentage correct) and reaction times are plotted for low load (blue bars) and high load (red bars). Error bars indicate the standard error of the sample mean, corrected to reflect the within-subjects contrast. Individual data are plotted and connected by grey lines to illustrate change in performance for each participant between low and high load conditions.

### Visual task

A significant effect of load on performance in the visual task was observed (Figure 2B). Increased load led to lower accuracy (Percentage Correct: *Mean*: LL = 94.5%, HL = 83.7%; *SD*: LL = 10.7, HL = 11.9; *t*(19) = 7.8, *p* < .001. d’: *Mean:* LL = 4.3, HL = 2.5; *SD*: LL = 1.3, HL = 1.0; *t*(19) = 9.9, *p* < .001) and longer RTs (*Mean*: LL = 594ms HL = 1000ms; *SD*: LL = 128, HL = 184; *t*(19) = −10.5, *p*< .001) confirming that the load manipulation was successful.

### Auditory processing

#### Effect of perceptual load on figure ground segregation

To quantify the effect of load on figure-ground segregation, we calculated a spatial filter which best captured the figure-specific brain response within the aggregate MEG signal (see methods). Figure 3A shows this filter (inset) and the derived evoked responses, contrasting the responses to FP and FA signals in low load (LL, left) and high load (HL, right). The similarity between FA and FP responses calculated here and those recorded in Experiment 1 confirms that the analysis was successful at isolating the relevant responses from within the mixture of auditory and visual evoked activity. A clear sustained figure-related negativity is observed in the LL condition from 35-60ms and from 110ms until the end of the epoch. In contrast, under high load the figure-related negativity was only significant from 225-265 ms – high visual load apparently reduced the auditory system’s ability to distinguish between FP and FA scenes. This is confirmed explicitly by evaluating the interaction between FP/FA and HH/LL conditions: for each subject a difference time-series (LL (FP-FA) – HL (FP – FA)) was computed and subjected to bootstrap resampling. Figure 3B plots the resulting mean difference across subjects. This indicated that the amplitude of the figure-related negativity was significantly different between load conditions from 290-475 ms post stimulus onset. To understand whether this effect is driven by a load effect on FP trials, FA trials, or both, we compared HL and LL responses for FP and FA stimuli separately. This analysis demonstrated that load only had a significant effect on FP responses: under high load, FP responses were reduced during the period from ~200-350 ms post stimulus (green horizontal lines in Figure 3B). Load did not have a significant effect on responses to FA stimuli.

**FIGURE 3:**
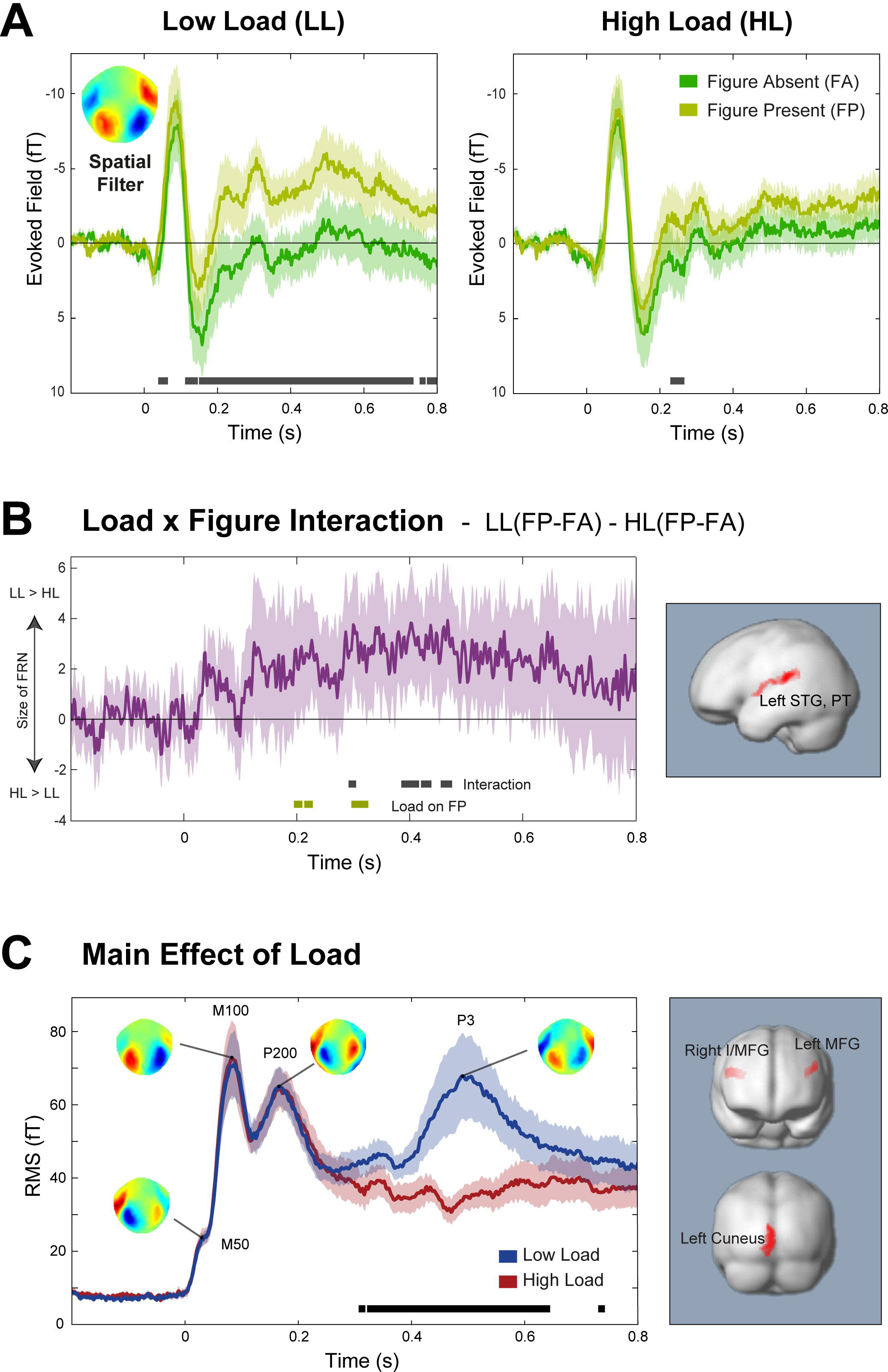
Experiment 2 - Effect of Load on Figure-Ground Segregation. **A. Figure Present/Absent responses as a function of visual load:** Evoked fields illustrating responses for Figure-present (FP) vs. Figure-absent (FA) signals under Low Load (LL; left) and High Load (HL; right) conditions. The spatial filter used to calculate the responses (see methods) is inset. Error bars reflect the SD of bootstrap resamplings for each condition, and significant differences between the conditions are marked by horizontal black bars at the bottom of the plots. **B. Interaction between load and figure present/absent**: The difference timeseries, LL(FP-FA) – HL(FP-FA), quantifies the interaction between load and figure. Error bars show 2*SD of the bootstrap, for comparison with zero line; periods when the values differed significantly from zero are indicated by the black bars below the plot. Green bars indicate periods when load had a significant effect on responses to FP stimuli (no significant periods were found for the effect of load on FA responses). Right panel indicates regions where the source analysis showed a significant interaction between responses in LL vs. HL and FP vs. FA. **C. Overall response to the SFG stimuli (collapsed over FA/FP conditions) as a function of load.** Mean RMS (instantaneous power) of responses to auditory stimuli (collapsed over FP and FA trials) in Experiment 2 under LL and HL, with scalp maps of peak topographies. The topographies are characteristic of auditory activity (symmetric dipolar pattern over temporal sensors) confirming that the source separation analysis was successful at isolating auditory activity. Error bars reflect the SD of bootstrap iterations for each condition, and significant differences between the conditions are marked by horizontal black bars at the bottom of the plot. Right panel shows regions where activity was stronger under LL than HL. No regions were found to be significant in the opposite direction to that displayed.

Overall, the data indicate that perceptual load had a significant effect on processing specifically related to detecting the figure. This pattern of results was replicated in a different experiment (see supplementary materials) which used the same SFG stimuli but a different load-inducing visual task.

**Table 2:** Experiment 2, Main Effect of Load. Source estimates for the difference between auditory responses in low and high load trials (regardless of figure presence).

| Cluster | *p*(FWE-corr) | Peaks |  | Co-ordinates (x,y,z) |  |  |
| --- | --- | --- | --- | --- | --- | --- |
|  |  | *t* | *p*(uncorr) |  |  |  |
| Cortical Structures |  |  |  |  |  |  |
| **Left Occipital Lobe:** | 0.001 | 4.46 | <.001 | -4 | -92 | -12 |
| Cuneus, Precuneus, Inferior Occipital Gyrus, BA 17, BA 18 |  | 3.92 | <.001 | -6 | -84 | 18 |
|  |  | 3.70 | <.001 | -2 | -100 | 2 |
| **Right Frontal Lobe:** | 0.025 | 3.96 | <.001 | 48 | 16 | 30 |
| Middle Frontal Gyrus, Inferior Frontal Gyrus |  | 3.94 | <.001 | 44 | 20 | 22 |
| **Left Frontal Lobe:** | 0.022 | 4.46 | <.001 | -48 | 20 | 30 |
| Middle Frontal Gyrus |  |  |  |  |  |  |

#### Source Analysis

When collapsed over low and high load, source analysis identified areas in the right temporal and right parietal lobes which showed greater activity in response to FP vs. FA scenes (Table 3). The temporal region covered the posterior portion of the right superior temporal gyrus, with some extension to middle temporal gyrus and planum temporale. This closely mirrors the bilateral temporal sources seen in Experiment 1, and previous fMRI and MEG data^25, 26^. The parietal source covered regions of the superior and inferior parietal lobules. It was slightly superior and anterior to the source seen in Experiment 1, and overall more diffuse, but given the relatively poor spatial resolution for MEG, we believe both represent activity within the IPS. Both these loci are also consistent with the fMRI and MEG data discussed above^25, 26^. This further confirms that the DSS analysis successfully captured the relevant SFG evoked activity.

**Table 3:**
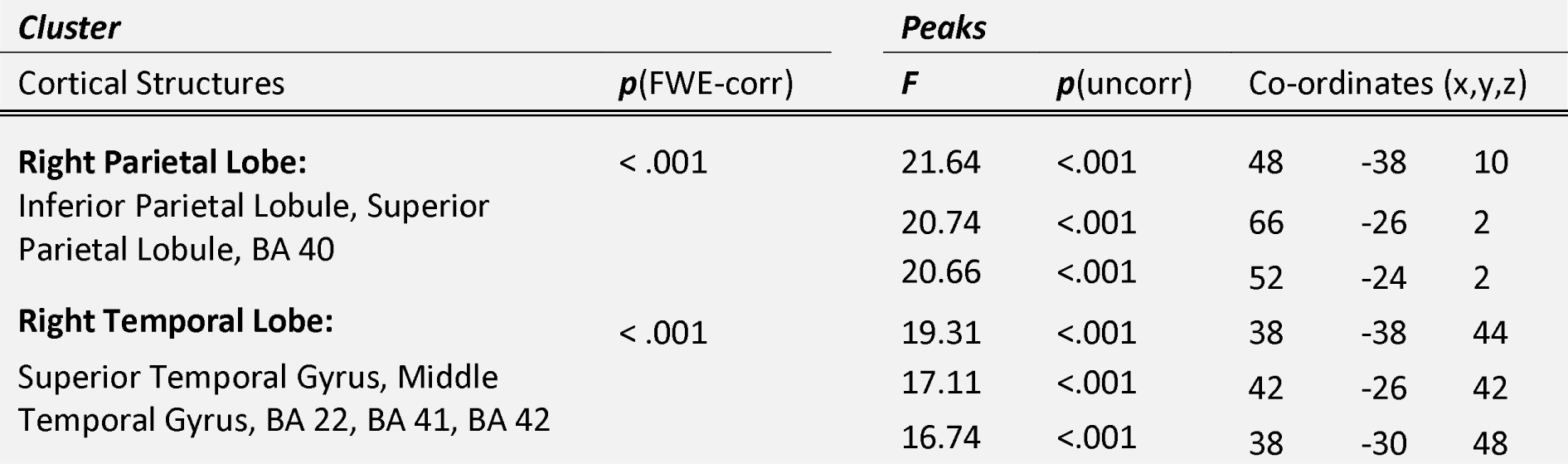
Experiment 2, Main Effect of Figure. Source estimates for the difference between auditory responses in FP and FA conditions (regardless of load).

The source of the interaction between load and figure was localised to the left temporal lobe (Figure 3B, Table 4). The area extends down the left superior temporal gyrus and planum temporale including Heschl’s gyrus. The data show that the successful differentiation of responses for FP vs. FA stimuli in this area is impaired under high load. This is commensurate with the main effect of figure described above, which was significant in the right but not left temporal gyrus. It suggests that under low load the SFG figures are processed in bilateral STG and right IPS (consistent with the data from Experiment 2), but as resources are depleted, processing in the left hemisphere is impaired more than that in the right.

**Table 4:**
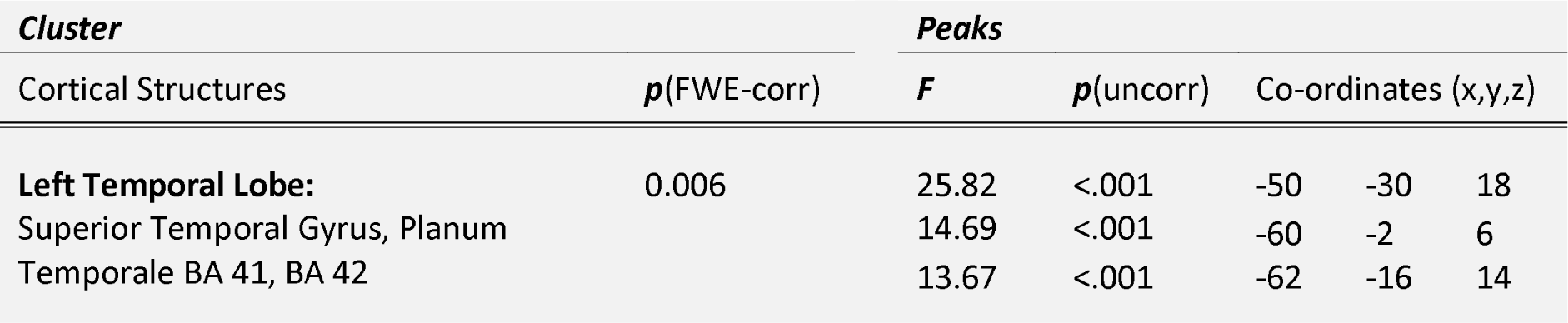
Experiment 2, Figure x Load Interaction. Source estimates for the interaction between load (low, high) and figure presence (FP, FA).

| *Cluster* |  | *Peaks* |  |  |  |  |
| --- | --- | --- | --- | --- | --- | --- |
| Cortical Structures | *p*(FWE-corr) | *F* | *p*(uncorr) | Co-ordinates (x,y,z) |  |  |
| **Left Temporal Lobe:** | 0.006 | 25.82 | <.001 | -50 | -30 | 18 |
| Superior Temporal Gyrus, Planum |  | 14.69 | <.001 | -60 | -2 | 6 |
| Temporale BA 41, BA 42 |  | 13.67 | <.001 | -62 | -16 | 14 |

#### Effect of perceptual load on overall response to ignored sound

To establish whether perceptual load had an effect on the base response to auditory stimuli (i.e. independently of whether a figure was present or absent), auditory components of the evoked response (after separation from visual evoked activity, but before isolating the figure response – see methods) were calculated. The responses (collapsed over FP and FA conditions) are illustrated in Figure 3C using the RMS over channels. The activity is characterized by the standard succession of auditory response peaks, and the field-maps associated with the major peaks (also plotted) exhibit the standard dipolar pattern over temporal channels commonly associated with auditory activity. The data closely match the responses observed during passive listening (Experiment 1, Figure 1B), confirming that the auditory activity was isolated successfully from the response mixtures.

Comparing responses under low and high visual perceptual load revealed significant effects of load from 300-650 ms post stimulus, with a clear P3 apparent in the responses to the sounds under low but not high visual load. This is in keeping with Molloy et al.’s previous findings^35^ where high visual load also led to the suppression of the P3 ‘awareness’ response to ignored sounds, and fits with a limited resource model. Molloy et al.^35^ also observed suppression of early (~100 ms) auditory responses under high load. In contrast, the present data show no evidence of a suppression of the onset responses under high visual load. This is likely because Molloy et al.^35^ used faint sounds whereas those used here were substantially louder.

Source analysis revealed significantly stronger activity in frontal and occipital regions in low load compared to high (see Figure 3A, Table 2). The activity in bilateral middle frontal gyri is likely to be the source of the P3 response which was apparent under low but not high load; the P3 has reliably been shown to have a generator in the frontal lobe when it occurs in response to non-target stimuli^36–39^. The difference in activity within the left occipital lobe may indicate some residual visual activity.

### Experiment 3: Psychophysics Dual Task

Experiment 3 was designed to determine whether the reduced figure-ground segregation that was shown in Experiment 2 under high (relative to low) visual load was associated with a reduction in perception of the auditory figures.

Results are shown in Table 5. Similarly to the behavioural pattern in Experiment 2, participants showed a significant effect of load on performance in the primary, visual, task. Increased load led to lower accuracy (*t*(1,11)=7.0, *p*<.001) and longer RTs (*t*(1,11)=-12.5, *p*<.001), indicating that the load manipulation was successful. Higher load in the primary visual task also led to poorer performance in the secondary, auditory, task: hit rates were reduced (t(1,11)=4.2, p =.001), and participants showed poorer sensitivity to the auditory target – d’ scores were significantly reduced in the high compared to the low load condition (*t*(1,11)=3.7, *p* =.004), with no change in decision criterion (beta) or false alarm rates.

**Table 5:** Experiment 3, Dual Task Behavioural Data.

|  | *Auditory Task* |  |  |  |  |  |  |  | *Visual Task* |  |  |  |
| --- | --- | --- | --- | --- | --- | --- | --- | --- | --- | --- | --- | --- |
|  | *d'* |  | *Beta* |  | *FA Rate* |  | *Hit Rate* |  | *% Correct* |  | *RT (ms)* |  |
|  | LL | HL | LL | HL | LL | HL | LL | HL | LL | HL | LL | HL |
| **Mean** | 1.8 | 1.5 | 0.80 | 0.91 | 31.1 | 32.8 | 85.5 | 80.5 | 97.5 | 89.9 | 714 | 1066 |
| **SD** | 0.27 | 0.23 | 0.16 | 0.15 | 5.7 | 5.9 | 3.3 | 3.6 | 0.5 | 1.4 | 42 | 31 |
| **p value** | 0.004 |  | 0.178 |  | 0.440 |  | 0.001 |  | <0.001 |  | <0.001 |  |

These data confirm that participants can successfully identify the figure within very short auditory scenes when these receive attentional resources in conditions of low perceptual load. However, high demands in a concurrent visual task reduced participants’ ability to segregate the auditory figures from the scene, leading to reduced detection sensitivity. This suggests that the reduction of the figure-related negativity seen under high visual load in the MEG data does indeed reflect a failure of the auditory system to segregate and thus hear an auditory figure out from a complex scene.

## Discussion

The present results revealed an extensive impact of visual attention on auditory figure-ground segregation. In conditions of low visual perceptual load, a neural marker of auditory figure-ground segregation (the figure-related negativity, FRN) was clearly evident from ~100-800 ms post sound onset. In contrast, conditions of high perceptual load in the visual task resulted in almost complete elimination of the neural response to the SFG figure, indicating a failure of auditory segregation in the absence of attention. Behavioural results also demonstrated poorer detection of the figure in conditions of high (vs. low) visual perceptual load even though participants were intently listening for it in both load conditions.

In addition to the effect of load on figure-ground segregation, high visual load also led to a reduced P3 response to sounds overall (irrespective of whether a figure was present). The P3 ‘awareness positivity’ is understood to reflect attentional engagement and conscious awareness of a stimulus^36, 38, 40, 41^. The finding that the auditory P3 is reduced under high visual load is consistent with a recent report that assessed responses to pure tones^35^ and confirms that under high visual load, unattended sounds are less likely to attract attention.

We note that with respect to overall response to the auditory stimuli (i.e. irrespective of figure vs. ground), the present findings were confined to a modulation of the P3 while previous work reported suppression of early evoked responses in auditory cortex^35, 42^. Molloy et al.^35^, for example, demonstrated that in addition to abolishing the P3 ‘awareness’ response, a high load visual task led to a reduction of the M100 onset response to concurrently presented pure tone stimuli. This likely occurred because Molloy et al. used faint sounds (12 dB above individual threshold). In contrast, the signals used in the present study were deliberately set loud enough to compensate for these early effects. Under these conditions, the effect of load was confined specifically to responses to figure-present stimuli, suggesting that visual perceptual load interfered with the processes which underlie the extraction of the figure and elicitation of the figure-related-negativity (FRN). Thus, our data indicate that apart from early effects on faint sounds, load on visual attention can also reduce the computational capacity of the auditory system.

The extent to which auditory figure-ground segregation is dependent on attention has been a longstanding issue in hearing research^4, 7–11, 30, 42–48^. Evidence of segregation in the supposed absence of directed attention (e.g.^30, 42, 47^) has been taken to indicate that segregation takes place automatically and pre-attentively, and that attention operates at a subsequent stage to enhance the representation of attended objects and suppress distracters^20, 49–52^, for example by modulating the temporal coherence of neural populations^3^. Specifically in the context of SFG segregation, accumulating work in humans has shown ‘figure’-evoked brain responses in naïve, distracted listeners, suggesting an early pre-attentive computation of temporal coherence^24–26^ that is enhanced during active listening^24^. In keeping with these reports, in the passive listening (Experiment 1) and low load (Experiment 2) conditions, the auditory evoked fields reliably showed a characteristic negativity in response to scenes where the figure was present relative to the figure-absent conditions, which we refer to as the figure-related negativity (FRN). However, we also show that depleting resources can lead to an abrupt failure of the process: conditions of high load in the visual task resulted in a reduction of the FRN so that there was almost no distinction between the responses to figure present and figure absent scenes. These results suggest that high load impairs the mechanisms which support auditory segregation, indicating that these computations are not performed in a fully automatic manner in the sense that they do require attentional resources.

Overall this pattern of results, where a process occurs when resources are available but can be disrupted in situations of very high demand, is congruent with load theory of attention (e.g.^27, 53^) and may explain some of the apparent discrepancies in previous research. Paradigms which attempted to direct attention away from the streaming signals but still saw evidence of successful segregation may have used distraction tasks which did not fully deplete resources. According to load theory, perceptual processing depends on limited resources that are allocated involuntarily to all stimuli within capacity. The level of perceptual load in the attended task therefore determines the extent to which any resources are available to ‘spill over’ to processing outside the focus of attention. The spill-over of attention allows figure-ground segregation in conditions of low perceptual load, while the greater level of engagement of resources in conditions of high perceptual load results in a significant decrease of both neural response to the figure and its conscious perception.

The FRN characterised here is generally reminiscent of the object-related negativity (ORN^29, 54, 55^, see also^33^) - a response from ~150-300 ms which is understood to be an index of segregation of a tone from a concurrent sound complex. Whilst both the FRN and ORN are associated with a similar perceptual outcome – the representation of a segregated ‘figure’ against a ‘background’ – it is likely that they arise from distinct underlying processes^23^. The calculations which underlie segregation based on harmonic cues presumably occur subcortically, during early stages of the auditory processing hierarchy^56^. In contrast, temporal coherence computation requires integration over time and likely has, at least in part, a cortical origin^25, 26, 46^. This difference may also make the process of SFG segregation more susceptible to depletion of resources than segregation based on harmonicity, consistent with previous studies which have reported no effect of attentional load on the ORN^30, 42^.

The neural underpinning for temporal coherence computations remain poorly understood and it is likely that the relevant computations are elaborated along a broad processing hierarchy. Current computational models of auditory scene analysis apply cross-channel correlation at a variety of temporal delays^2, 20, 21^ that are hypothesized to be supported by rapid adaptive processes in auditory cortex^46^. Recent animal work has indicated that this rapid plasticity only takes place when animals are explicitly attending to the auditory signals^46^, which does not tally with the human literature, where segregation is reliably observed during passive listening^24, 26^. However, our data shows that the limiting aspect is not active attention per se, but rather the availability of general computational resources irrespective of the specific focus of attention. Thus, a disparity in the size of resource pools between humans and animals could lead to the apparent differences between the levels of attention required for these adaptive processes.

Human neuroimaging work has implicated Planum Temporale (PT) and Intra-Parietal Sulcus (IPS) in the process of detecting SFG ‘figures’ ^25, 26^ and a similar network is also seen in the present experiments during low load/passive listening conditions (Experiment 1 here). The involvement of PT in segregation is also in agreement with other work using simpler streaming signals^57–59^. Teki et al.^25^ suggest that PT operates as a hub for the process that computes the coherence maps, while IPS is involved in reading out these calculations and encoding the signal as consisting of several sources.

Interestingly, here load specifically impacted processing in temporal cortex: the effect of load on the FRN was localised to the upper bank of the left superior temporal gyrus, including the PT and Heschl’s gyrus. Activity in this region was generally more pronounced during figure present scenes compared to figure absent, but the distinction was less marked under high load. This is consistent with both the findings of attention-dependent adaptation in A1^46^, and with the hypothesized role of PT in computing the temporal coherence maps which underlie segregation^25^. Importantly, our data demonstrate that processing within these regions is not encapsulated but draws on domain-general resources such that conditions of high demand in the visual modality can lead to the failure of fundamental aspects of auditory processing.

## Materials & Methods

### Experiment 1: MEG to short SFG sequences

#### Participants

Sixteen paid participants (9 male; mean age of 24.8 years, SD = 3.0 years) took part in Experiment 1. All were right handed, had normal or corrected to normal vision and reported normal hearing and no history of neurological disorders. The experimental protocol for all reported experiments was approved by the University College London research ethics committee.

#### Apparatus, Stimuli and Procedure

The magnetic signals were recorded using a CTF-275 MEG system (axial gradiometers, 274 channels, 30 reference channels, VSM MedTech, Canada) in a magnetically shielded room. Subjects were seated in an upright position, with the visual stimuli projected onto a screen placed ~52cm from the participants’ eyes. Data were recorded continuously with a 600Hz sampling rate and a 100Hz hardware low-pass filter.

The auditory stimuli were ~200ms long, diotically presented SFG stimuli ^2, 25, 26^ (see Figure 1). Signals consisted of a succession of chords, each comprised of multiple frequency components. Frequencies were chosen from a log-distributed pool of 129 frequencies from 179 to 7246 Hz. Each chord was comprised of between 11 and 21 (number was uniformly distributed) frequency components, which were selected from the frequency pool with equal probability. The *‘figure-absent’* stimuli (FA, 50%) were formed of random frequency chords. The *‘figure-present’* stimuli (FP, 50%) were constrained so that a subset of the frequencies were repeated in each chord (this parameter is referred to as the ‘coherence’ of the figure, see Teki et al., 2011; 2013; 2016), while the others were selected randomly for each chord. The repetition of the coherent frequencies creates the auditory ‘figure’ which can be heard separately from the stochastically changing background^2^.

The stimuli were varied along several parameters so as to optimize the stimuli to be used for the investigation of load (Experiment 2). Specifically, we varied the number of coherent frequencies used for the figure (6 or 8); the duration of the chords (25 or 30ms); and the number of chords (6 or 8); all combinations were tested (8 possibilities) and stimuli were either FP or FA, creating 16 conditions. 120 exemplars of each of the 16 conditions were randomly allocated into 4 blocks of 480 stimuli, and presented with ISIs of 800ms. Naïve participants passively listened to the signals while performing an incidental visual task. Prior to the recording, the volume of the stimuli was set to a comfortable level (~70 dB SPL) by each participant.

The visual task was designed to be very low demand. Pictures of landscapes were presented in groups of three (5 seconds per image, fade in and out over 1 second), and participants had to press a button if picture 2 or 3 was the same as picture 1. Instances of repetitions were relatively rare (~1 in 12 sets) so that motor responses were kept to a minimum. This task helped ensure that participants’ eyes were open and they were awake throughout the blocks, but did not place a high demand on processing resources.

At the beginning of the session, a short (4 minutes) ‘localizer’ block was recorded in order to characterise participants’ neural responses to simple auditory stimuli. The measurement consisted of 200 presentations of a 100ms long, 1 kHz pure tone with ISIs randomly distributed between 700 and 1500ms. Participants watched a static fixation cross in the centre of the screen and were not required to perform a task.

#### Analysis

All conditions (over coherence, chord duration and number of chords) showed very similar evoked responses; the data were therefore collapsed over all conditions for display purposes in the results, and for the subsequent source analysis.

The data from the localizer block were divided into 800ms epochs, and baseline-corrected using a 200ms pre-stimulus interval. The M100 onset response (Roberts et al., 2000) was identified for each subject as a source/sink pair in the magnetic-field contour plots distributed over the temporal region of each hemisphere. For each subject, the 40 most activated channels at the peak of the M100 (20 in each hemisphere) were selected for subsequent sensor-level analysis of the responses evoked by the SFG stimuli.

The data from the main blocks were epoched into 1000ms trials which covered 800ms post-stimulus onset, and 200ms pre-onset. All data were baseline corrected to the pre-onset interval. Epochs with amplitudes above 3pT (~6% of trials) were considered to contain artefacts and discarded. A PCA-based, Denoising Source Separation (DSS; ^60^) routine was applied to the data to extract stimulus-locked activity. The 20 most repeatable components were retained and projected back to sensor space. To characterize the response at this stage, the root mean square (RMS) of the evoked field over the localizer channels was calculated for each time point to give a time-series which reflects the instantaneous power of the evoked response. For illustrative purposes, group-RMS (RMS of individual subject RMSs) is plotted (Figure 1B) but statistical analysis was performed across subjects.

To characterise the elements of the response which are specific to FP stimuli, a further DSS was conducted, this time optimized to find components (spatial filters) which differed maximally between FA and FP trials (calculated over all channels and the whole epoch; ^60^). The highest ranked DSS component was retained for the analysis and used as a spatial filter (source model) for the analysis of FA vs. FP trials (Figure 1C). In all cases this spatial filter corresponded to the standard temporal dipolar pattern associated with auditory responses.

FP trials were characterized by increased negativity relative to FA trials. To quantify this effect, the difference between the evoked responses for FP and FA trials was calculated for each participant, and subjected to bootstrap re-sampling (1000 iterations, balanced; ^61^). The difference was judged to be significant if the proportion of bootstrap iterations which fell above/below zero was more than 99% (i.e. *p*<.01) for 12 or more adjacent samples (20ms). The bootstrap analysis was run over the entire epoch duration (200ms pre onset to 800ms post onset); all significant intervals identified in this way are indicated in the figure.

Sources were estimated using multiple sparse priors (MSP; ^62^) analysis. Inversions were based on all MEG channels and used a single shell head model with group constraints. Second-level analyses consisted of t-contrasts to compare activation between FP and FA conditions. Results were thresholded at *p*< .001 at the peak level and then subjected to a whole brain *p*< .05 FWE correction at the cluster level. In one instance a small-volume correction (a 10mm diameter sphere centred at the peak of the cluster) was applied instead, since the cluster was small but in a location consistent with previous fMRI^26^ and MEG^25^ sources for similar SFG stimuli. The use of a different correction is marked in Table 2.

### Experiment 2: Effect of Visual Load on Figure-Ground Segregation

#### Participants

Twenty paid participants (8 male; mean age of 24.5 years, SD = 4.3 years) took part in Experiment 2. All were right handed, had normal or corrected to normal vision and reported normal hearing and no history of neurological disorders.

#### Apparatus and Stimuli

The apparatus and recording methods were identical to those in Experiment 1.

The visual search arrays, presented for 200 ms on a dark grey background, consisted of five coloured shapes spaced equally around a (non-visible) circle centred at fixation and subtending 1.9 viewing angle. The five shapes comprised one each of a circle, triangle, square, diamond and pentagon. The colours were assigned so that there were always two red items, two green, and one either blue or yellow (50% of trials each). In principle any display could be used as a low load (LL; colour search) or high load (HL; colour-shape combination search) stimulus, so that displays were identical between load conditions. The target in low load was any blue shape; the high load targets were a red circle or green square. Targets were present in 50% of arrays, and counterbalanced so that LL and HL targets did not correlate (i.e. if the LL target was present in an array, the likelihood of the HL target being present was 50%, and vice-versa). The positions of the shapes were pseudo-randomised on each trial so that the target had an equal probability of occurring in each position. Each combination of shape and colour was equiprobable across the stimulus array sets.

On half of the trials, the visual display was accompanied by a brief auditory stimulus, presented at the same time and for the same duration as the display (Figure 2). The auditory stimuli were identical to the SFG stimuli used in Experiment 1, but with fixed parameters: coherence 6, chord duration 25ms and chord number 8, producing a 200ms long stimulus (FP and FA with equal probability). There was no active auditory task - participants were encouraged to focus on the visual task and were told that the sounds were incidental.

The session comprised 12 blocks (6 low load, 6 high load) consisting of 80 trials each; the order of the blocks was counterbalanced between participants. Trial-by-trial feedback was not given, but at the end of each block, participants were provided a score of percentage correct on the visual task, to boost engagement. Blocks lasted for ~4 minutes each, and participants were encouraged to take breaks between blocks when needed.

#### Procedure

Figure 2 shows a schematic diagram of the trial structure. Each trial began with a fixation cross presented at the centre of the screen for 1000ms. Subsequently, a visual search array was presented for 200ms, accompanied on 50% of trials by an auditory stimulus. A blank screen was then presented for 1800ms, during which participants were to make a speeded response as to whether the visual target was present or absent (by pressing one of two buttons with their right hand).

#### Analysis

The data from the main blocks were epoched into 1000ms trials, including a 200ms pre-onset interval. All data were baseline corrected to the pre-onset interval. Epochs with amplitudes above 3pT (~6% of trials) were considered to contain artefacts and discarded. DSS^60^ was applied to the data to extract stimulus-locked activity. As with the previous analysis, the 20 most repeatable components were retained.

Since the auditory stimuli were always presented concurrently with the visual search array, a further DSS step was necessary to separate auditory responses from the measured auditory-visual combined response. This analysis (collapsed over load conditions) identified components in the data which showed the greatest difference between trials when the visual stimuli were presented alone (50%) and those when an auditory stimulus was also present (50%), with the aim of isolating activity which specifically relates to the auditory stimuli (see detailed description of the procedure in ^35, 60^). The ten highest ranked components were projected back into channel space and the dataset was split into the low and high load conditions. Note that the RMS and scalp topographies (Figure 3A) of the auditory component calculated from this analysis closely resemble the data recorded in response to the same stimuli in Experiment 1 (Figure 1B), demonstrating that the auditory evoked activity was successfully extracted.

As in Experiment 1, a subsequent DSS analysis was applied (collapsed over load conditions) to produce a spatial filter which reflects activity most different between FP and FA trials. The data were then separated into LL/HL and FP/FA conditions for analysis (Figure 3). Statistical analyses were as described for Experiment 1.

Source inversions were calculated using multiple sparse priors (MSP; ^62^) analysis. Inversions were based on all MEG channels and used a single shell head model and group constraints. For estimating sources of auditory activity, a soft prior over temporal and parietal areas was used, motivated by previous fMRI and MEG data for SFG stimuli^25, 26^, and our source results from Experiment 2, all of which indicate potential sources throughout the temporal lobe and in IPS. The prior mask was created in FSLview (http://surfer.nmr.mgh.harvard.edu/), based on combining the Harvard–Oxford Structural atlases for all temporal areas, and the Juelich histologic atlas for IPS, with a threshold of 5%. This resulted in a very broad prior, which was binarised so that the strength was equal over all regions. Note that solutions were not restricted to this mask, it served only as a prior to the source algorithm.

Second-level analyses consisted of paired t-contrasts to compare the visual and auditory responses between low and high load, and a full factorial RM F-contrast to model the auditory responses, including main effects of load and figure, and the load x figure interaction. Results were thresholded at *p*< .001 at the peak level and then subjected to a whole brain *p*< .05 FWE correction at the cluster level.

### Experiment 3: Psychophysics Dual Task

#### Participants

Thirteen paid participants took part in the behavioural study. One was excluded because their performance on the low-load task was extremely poor (61%; average of all included participants was 97.5%). For the remaining twelve participants (8 female), ages ranged from 18-35 years (mean = 21.4, SD = 4.1). All participants had normal or corrected to normal vision and reported normal hearing.

#### Apparatus and Stimuli

The experiment was run on a Dell PC with a 13” monitor using Matlab 7.12 and Cogent 2000 (http://www.vislab.ucl.ac.uk/cogent.php). A viewing distance of 57cm was maintained throughout using a chin rest. Sounds were presented via tubephones (E-A-RTONE 3A 10 Ω, Etymotic Research, Inc) inserted into the ear-canal.

The stimuli were identical to those used in Experiment 2.

#### Procedure

Trials were similar to those in Experiment 2, except that the auditory stimuli were present on every trial, and participants were asked to perform a dual task, responding first to the visual search (target present or absent), and subsequently to the presence of an SFG ‘figure’ (FP stimuli). Trials were similar to those in Experiment 2, but after the response to the visual search (using their right hand), a prompt was displayed on the screen for 2000ms (see Figure 2A) during which participants indicated whether they had heard the auditory figure by pressing a button with their left hand. The experiment consisted of 12 blocks of 40 trials each, six low load and six high, with the order of blocks counterbalanced between participants.

The experimental session was preceded by a series of short demo blocks (with trial-by-trial feedback) which introduced the auditory and visual tasks separately, and then combined them to illustrate the procedure for the dual task.

## ACKNOWLEDGEMENTS

We are grateful to the radiographer team at the UCL Wellcome Trust Centre for Neuroimaging for excellent MEG technical support. This study was supported by a Wellcome Trust project grant to MC (093292/Z/10/Z). KM is supported by an Economic and Social Research Council studentship (ES/J500185/1).

## References

1. Lavie N. Perceptual load as a necessary condition for selective attention. Journal of experimental psychology Human perception and performance 21, 451–468 (1995).

2. Teki S, Chait M, Kumar S, Shamma S, Griffiths TD. Segregation of complex acoustic scenes based on temporal coherence. eLife 2, e00699 (2013).

3. Shamma SA, Micheyl C. Behind the scenes of auditory perception. Current opinion in neurobiology 20, 361–366 (2010).

4. Shamma SA, Elhilali M, Micheyl C. Temporal coherence and attention in auditory scene analysis. Trends in neurosciences 34, 114–123 (2011).

5. Snyder JS, Gregg MK, Weintraub DM, Alain C. Attention, awareness, and the perception of auditory scenes. Frontiers in psychology 3, 15 (2012).

6. Puvvada KC, Simon JZ. Cortical Representations of Speech in a Multitalker Auditory Scene. The Journal of neuroscience: the official journal of the Society for Neuroscience 37, 9189–9196 (2017).

7. Carlyon RP, Cusack R, Foxton JM, Robertson IH. Effects of attention and unilateral neglect on auditory stream segregation. Journal of experimental psychology Human perception and performance 27, 115–127 (2001).

8. Cusack R, Deeks J, Aikman G, Carlyon RP. Effects of location, frequency region, and time course of selective attention on auditory scene analysis. Journal of experimental psychology Human perception and performance 30, 643–656 (2004).

9. Macken WJ, Tremblay S, Houghton RJ, Nicholls AP, Jones DM. Does auditory streaming require attention? Evidence from attentional selectivity in short-term memory. Journal of experimental psychology Human perception and performance 29, 43–51 (2003).

10. Thompson SK, Carlyon RP, Cusack R. An objective measurement of the build-up of auditory streaming and of its modulation by attention. Journal of experimental psychology Human perception and performance 37, 1253–1262 (2011).

11. Weintraub DM, Metzger BA, Snyder JS. Effects of attention to and awareness of preceding context tones on auditory streaming. Journal of experimental psychology Human perception and performance 40, 685–701 (2014).

12. Carlyon RP. How the brain separates sounds. Trends in cognitive sciences 8, 465–471 (2004).

13. Fishman YI, Steinschneider M. Neural correlates of auditory scene analysis based on inharmonicity in monkey primary auditory cortex. The Journal of neuroscience: the official journal of the Society for Neuroscience 30, 12480–12494 (2010).

14. Gutschalk A, Micheyl C, Oxenham AJ. Neural correlates of auditory perceptual awareness under informational masking. PLoS biology 6, e138 (2008).

15. Kidd G, Jr., Richards VM, Streeter T, Mason CR, Huang R. Contextual effects in the identification of nonspeech auditory patterns. The Journal of the Acoustical Society of America 130, 3926–3938 (2011).

16. Micheyl C, et al. The role of auditory cortex in the formation of auditory streams. Hearing research 229, 116–131 (2007).

17. Micheyl C, Oxenham AJ. Across-frequency pitch discrimination interference between complex tones containing resolved harmonics. The Journal of the Acoustical Society of America 121, 1621–1631 (2007).

18. Moore BC, Gockel HE. Properties of auditory stream formation. Philosophical transactions of the Royal Society of London Series B, Biological sciences 367, 919–931 (2012).

19. Fishman YI, Kim M, Steinschneider M. A Crucial Test of the Population Separation Model of Auditory Stream Segregation in Macaque Primary Auditory Cortex. The Journal of neuroscience: the official journal of the Society for Neuroscience 37, 10645–10655 (2017).

20. Elhilali M, Ma L, Micheyl C, Oxenham AJ, Shamma SA. Temporal coherence in the perceptual organization and cortical representation of auditory scenes. Neuron 61, 317–329 (2009).

21. Krishnan L, Elhilali M, Shamma S. Segregating complex sound sources through temporal coherence. PLoS computational biology 10, e1003985 (2014).

22. Micheyl C, Hanson C, Demany L, Shamma S, Oxenham AJ. Auditory stream segregation for alternating and synchronous tones. Journal of experimental psychology Human perception and performance 39, 1568–1580 (2013).

23. Micheyl C, Kreft H, Shamma S, Oxenham AJ. Temporal coherence versus harmonicity in auditory stream formation. The Journal of the Acoustical Society of America 133, EL188–194 (2013).

24. O’Sullivan JA, Shamma SA, Lalor EC. Evidence for Neural Computations of Temporal Coherence in an Auditory Scene and Their Enhancement during Active Listening. The Journal of neuroscience: the official journal of the Society for Neuroscience 35, 7256–7263 (2015).

25. Teki S, Barascud N, Picard S, Payne C, Griffiths TD, Chait M. Neural Correlates of Auditory Figure-Ground Segregation Based on Temporal Coherence. Cereb Cortex 26, 3669–3680 (2016).

26. Teki S, Chait M, Kumar S, von Kriegstein K, Griffiths TD. Brain bases for auditory stimulus-driven figure-ground segregation. The Journal of neuroscience: the official journal of the Society for Neuroscience 31, 164–171 (2011).

27. Lavie N. Distracted and confused?: selective attention under load. Trends in cognitive sciences 9, 75–82 (2005).

28. Lavie N, Beck DM, Konstantinou N. Blinded by the load: attention, awareness and the role of perceptual load. Philosophical transactions of the Royal Society of London Series B, Biological sciences 369, 20130205 (2014).

29. Alain C, Arnott SR, Picton TW. Bottom-up and top-down influences on auditory scene analysis: evidence from event-related brain potentials. Journal of experimental psychology Human perception and performance 27, 1072–1089 (2001).

30. Alain C, Izenberg A. Effects of attentional load on auditory scene analysis. Journal of cognitive neuroscience 15, 1063–1073 (2003).

31. Alain C, McDonald KL. Age-related differences in neuromagnetic brain activity underlying concurrent sound perception. The Journal of neuroscience: the official journal of the Society for Neuroscience 27, 1308–1314 (2007).

32. McDonald KL, Alain C. Contribution of harmonicity and location to auditory object formation in free field: evidence from event-related brain potentials. The Journal of the Acoustical Society of America 118, 1593–1604 (2005).

33. Toth B, Kocsis Z, Haden GP, Szerafin A, Shinn-Cunningham BG, Winkler I. EEG signatures accompanying auditory figure-ground segregation. NeuroImage 141, 108–119 (2016).

34. Treisman AM, Gelade G. A feature-integration theory of attention. Cognitive psychology 12, 97–136 (1980).

35. Molloy K, Griffiths TD, Chait M, Lavie N. Inattentional Deafness: Visual Load Leads to Time-Specific Suppression of Auditory Evoked Responses. The Journal of neuroscience: the official journal of the Society for Neuroscience 35, 16046–16054 (2015).

36. Comerchero MD, Polich J. P3a and P3b from typical auditory and visual stimuli. Clinical neurophysiology: official journal of the International Federation of Clinical Neurophysiology 110, 24–30 (1999).

37. Goldstein A, Spencer KM, Donchin E. The influence of stimulus deviance and novelty on the P300 and novelty P3. Psychophysiology 39, 781–790 (2002).

38. Polich J. Updating P300: an integrative theory of P3a and P3b. Clinical neurophysiology: official journal of the International Federation of Clinical Neurophysiology 118, 2128–2148 (2007).

39. Simons RF, Graham FK, Miles MA, Chen X. On the relationship of P3a and the Novelty-P3. Biological psychology 56, 207–218 (2001).

40. Kok A. On the utility of P3 amplitude as a measure of processing capacity. Psychophysiology 38, 557–577 (2001).

41. Picton TW. The P300 wave of the human event-related potential. Journal of clinical neurophysiology: official publication of the American Electroencephalographic Society 9, 456–479 (1992).

42. Dyson BJ, Alain C, He Y. Effects of visual attentional load on low-level auditory scene analysis. Cognitive, affective & behavioral neuroscience 5, 319–338 (2005).

43. Alain C, Arnott SR. Selectively attending to auditory objects. Frontiers in bioscience: a journal and virtual library 5, D202–212 (2000).

44. Dyson BJ, Alain C. Representation of concurrent acoustic objects in primary auditory cortex. The Journal of the Acoustical Society of America 115, 280–288 (2004).

45. Lipp R, Kitterick P, Summerfield Q, Bailey PJ, Paul-Jordanov I. Concurrent sound segregation based on inharmonicity and onset asynchrony. Neuropsychologia 48, 1417–1425 (2010).

46. Lu K, Xu YB, Yin PB, Oxenham AJ, Fritz JB, Shamma SA. Temporal coherence structure rapidly shapes neuronal interactions. Nat Commun 8, (2017).

47. Masutomi K, Barascud N, Kashino M, McDermott JH, Chait M. Sound Segregation via Embedded Repetition Is Robust to Inattention. J Exp Psychol Human 42, 386–400 (2016).

48. Sussman ES, Horvath J, Winkler I, Orr M. The role of attention in the formation of auditory streams. Perception & psychophysics 69, 136–152 (2007).

49. Akram S, Englitz B, Elhilali M, Simon JZ, Shamma SA. Investigating the neural correlates of a streaming percept in an informational-masking paradigm. PloS one 9, e114427 (2014).

50. Bidet-Caulet A, Fischer C, Besle J, Aguera PE, Giard MH, Bertrand O. Effects of selective attention on the electrophysiological representation of concurrent sounds in the human auditory cortex. The Journal of neuroscience: the official journal of the Society for Neuroscience 27, 9252–9261 (2007).

51. Ding N, Simon JZ. Emergence of neural encoding of auditory objects while listening to competing speakers. Proceedings of the National Academy of Sciences of the United States of America 109, 11854–11859 (2012).

52. Zion Golumbic EM, et al. Mechanisms underlying selective neuronal tracking of attended speech at a "cocktail party". Neuron 77, 980–991 (2013).

53. Lavie N. Attention, Distraction, and Cognitive Control Under Load. Curr Dir Psychol Sci 19, 143–148 (2010).

54. Alain C, Schuler BM, McDonald KL. Neural activity associated with distinguishing concurrent auditory objects. The Journal of the Acoustical Society of America 111, 990–995 (2002).

55. Alain C, Theunissen EL, Chevalier H, Batty M, Taylor MJ. Developmental changes in distinguishing concurrent auditory objects. Brain research Cognitive brain research 16, 210–218 (2003).

56. Micheyl C, Oxenham AJ. Pitch, harmonicity and concurrent sound segregation: psychoacoustical and neurophysiological findings. Hearing research 266, 36–51 (2010).

57. Gutschalk A, Oxenham AJ, Micheyl C, Wilson EC, Melcher JR. Human cortical activity during streaming without spectral cues suggests a general neural substrate for auditory stream segregation. The Journal of neuroscience: the official journal of the Society for Neuroscience 27, 13074–13081 (2007).

58. Schadwinkel S, Gutschalk A. Activity associated with stream segregation in human auditory cortex is similar for spatial and pitch cues. Cereb Cortex 20, 2863–2873 (2010).

59. Wilson EC, Melcher JR, Micheyl C, Gutschalk A, Oxenham AJ. Cortical FMRI activation to sequences of tones alternating in frequency: relationship to perceived rate and streaming. Journal of neurophysiology 97, 2230–2238 (2007).

60. de Cheveigné A, Parra LC. Joint decorrelation, a versatile tool for multichannel data analysis. NeuroImage 98, 487–505 (2014).

61. Efron B, Tibshirani RJ. An Introduction to the Bootstrap: Monographs on Statistics and Applied Probability, Vol. 57. New York and London: Chapman and Hall/CRC, (1993).

62. Litvak V, Friston K. Electromagnetic source reconstruction for group studies. NeuroImage 42, 1490–1498 (2008).

63. Moore BC, Glasberg BR. Suggested formulae for calculating auditory-filter bandwidths and excitation patterns. The Journal of the Acoustical Society of America 74, 750–753 (1983).

64. Plack CJ, Moore BC. Temporal window shape as a function of frequency and level. The Journal of the Acoustical Society of America 87, 2178–2187 (1990).

